# Ultra-widefield fluorescein angiography findings in patients with multiple retinal holes

**DOI:** 10.1101/578328

**Authors:** Jiwon Baek, Byungju Jung, Hyunggu Kwon, Sohee Jeon

## Abstract

**Purpose:** To evaluate the ultra-widefield fluorescein angiography (UWFA) findings in patients with multiple retinal holes.

**Methods:** Seventy-five eyes, each with more than two retinal holes, underwent a comprehensive ophthalmologic examination, including UWFA, along with 77 age-matched control eyes with no retinal holes. The UWFA was scored according to a system suggested by the Angiography Scoring for Uveitis Working Group (ASUWG). UWFA findings from the patients with retinal holes were compared with those of the control group without retinal holes. Factors associated with a high ASUWG score were also evaluated.

**Results:** Patients with multiple retinal holes showed a significantly higher prevalence of retinal vascular staining/leakage, capillary leakage at the posterior pole, and capillary leakage at the periphery when compared to the control group (*p*<0.001, for each of them). Univariate analysis revealed that logMAR BCVA (r=0.271, *p*=0.027), spherical equivalent (r= −0.275, *p*=0.021), and number of retinal holes (r=0.271, p=0.027) were associated with a higher ASUWG score. After adjustments for age, gender, and logMAR BCVA, multivariate regression analysis revealed that the spherical equivalent was independently associated with a higher ASUWG score (r^2^=0.161, *p*=0.001).

**Conclusions:** Patients with multiple retinal holes showed profound peripheral vascular leakages on UWFA findings, suggesting the presence of chronic retinal traction induced by equatorial scleral elongation.

---

Rhegmatogenous retinal detachment (RRD) is a potentially devastating condition which arise from posterior vitreous detachment (PVD) and retinal breaks.^1,2^ Retinal breaks are classified into retinal tears or holes according to the morphology and its relationship with the vitreous.^3^ Retinal tears develop when there is a strong attachment of the retina to the vitreous, producing traction on susceptible areas of retina during the process of separation from vitreous from retina, and eventually causing RRD when the fluid enters the subretinal space through the tears. However, the association between retinal holes and PVD has not yet been clearly elucidated. Retinal holes are not usually associated with visible vitreous traction, and retinal hole-associated RRDs may be found without any obvious PVD.^4,5^ Although some forms of viral retinitis, such as acute retinal necrosis and cytomegalovirus retinitis, have reportedly resulted in multiple retinal holes at the junction between normal and necrotic retina,^6^ there has been limited data to date on the cause of retinal holes in eyes without retinal inflammation.

We hypothesized that patients with multiple retinal holes might in fact have undiagnosed peripheral retinal pathologies that might not easily be detected by conventional methods. The use of ultra-widefield fluorescein angiography (UWFA) has enabled us to better understand the pathological features of various peripheral retinal diseases.^7,8^ We speculated that UWFA imaging would help facilitate an understanding of the pathophysiology of multiple retinal holes. In this study, therefore, we evaluated the UWFA findings of patients with multiple retinal holes and compared them with those of normal controls.

## Methods

The case group in this multicenter retrospective study included patients who visited Seoul St. Mary’s Hospital, Bucheon St. Mary’s Hospital, and the Keye Eye Center for multiple retinal holes, between March 1, 2016, and June 28, 2018. The control group consisted of the other eyes of patients who visited these same institutions with central serous chorioretinopathy and branched retinal vein occlusion over same period. Institutional Review Board (IRB)/Ethics Committee approval was obtained for the study. The study protocol adhered to the principles of the Declaration of Helsinki.

The diagnosis of retinal hole was made by the physician performing the fundus examination, when a retinal break was round-shaped, and not associated with vitreoretinal traction.^3^ The exclusion criteria were as follows: (1) history of other retinal diseases, such as age-related macular degeneration or diabetic retinopathy; (2) media opacity that markedly obscured the image on optical coherence tomography (OCT) or UWFA; (3) history of intraocular surgery including cataract surgery; (4) ocular trauma; (5) presence of intraocular inflammatory changes at presentation; and (6) history of uveitis.

Comprehensive ocular examinations were performed, including best corrected visual acuity (BCVA), intraocular pressure (IOP) measurement, slit-lamp examination, refractive error, ultra-widefield fundus photography, and UWFA (Optos Optomap Panoramic 200A Imaging System; Optos plc, Dunfermline, Scotland). UWFA images were obtained after intravenous injection of 10% fluorescein (Fluorescite 10%; Alcon, Ft. Worth, TX, USA). We evaluated UWFA images according to the original scoring system suggested by the Angiography Scoring for Uveitis Working Group (ASUWG).^9^ The following findings were recorded: optic disc hyperfluorescence detected at 5–10 min, macular edema at 10 min, retinal vascular staining or leakage, capillary leakage at the posterior pole or periphery, retinal capillary nonperfusion, neovascularization, pinpoint leakage, retinal staining or subretinal pooling at 5–10 min (Fig 1). Two investigators (JB, BJ) measured the UWFA parameters, and when the results were discordant, a third investigator (SJ) was invited to give an additional opinion. All investigators were blinded to the clinical data and undertook the scoring independently.

**Figure 1.**
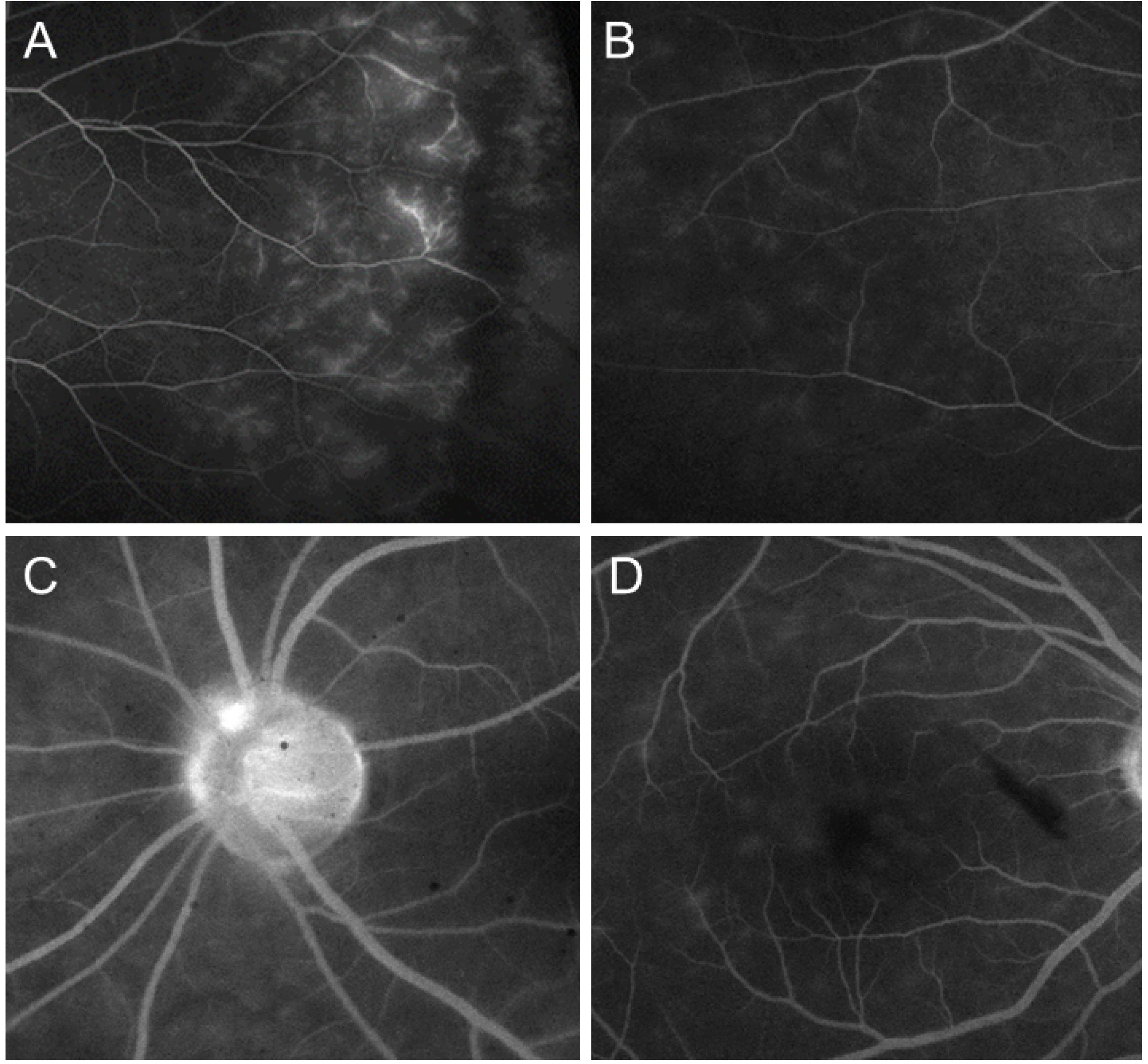
Representative images of enrolled patients who showed positive results for UWFA imaging. Peripheral retinal vascular leakages (A), peripheral capillary staining or leakages (B), optic disc staining (C), and posterior pole capillary leakages (D) are seen.

## Statistical analysis

SPSS, version 15.0 for Windows (SPSS, Inc., Chicago, IL, USA) was used for statistical analysis. Descriptive data were recorded as means ± standard deviation unless otherwise specified. Chi-square and two-tailed t-tests were used to assess the differences between patients with and without multiple retinal holes. To understand more about the association between the clinical data and the presence of retinal vascular leakage, we undertook additional analysis of the ASUWG scores. The Shapiro-Wilk test was used to assess normality. The Pearson correlation coefficient or Spearman rank correlation coefficient was determined to assess the association between continuous variables, according to the normality of distribution. Independent variables significantly associated with ASUWG scores in univariate analysis (*p*<0.05) and potentially confounding parameters were included as independent covariables in multivariate analysis by multiple regression analysis. A *p* value <0.05 was considered statistically significant.

## Results

The medical records of a total of 75 eyes with multiple retinal holes, and 77 control eyes, were reviewed. The baseline demographics are summarized in Table 1. There was no significant difference in the age, gender, the presence of diabetes or hypertension between two groups (p=0.222, 0.332, 0.497, and 0.746, respectively). The presence of other systemic diseases was not reported at the time of the initial visit in both groups. There was no significant difference in the mean LogMAR BCVA (*p*=0.265) and IOP (*p*=0.091). The mean refractive error was −5.01 ± 3.38 diopter in the case group, and −1.72 ± 2.49 in the control group (*p*<0.001). The mean number of retinal holes per eye was 3.26±1.53 (range, 2–9) in the study group.

**Table 1.**
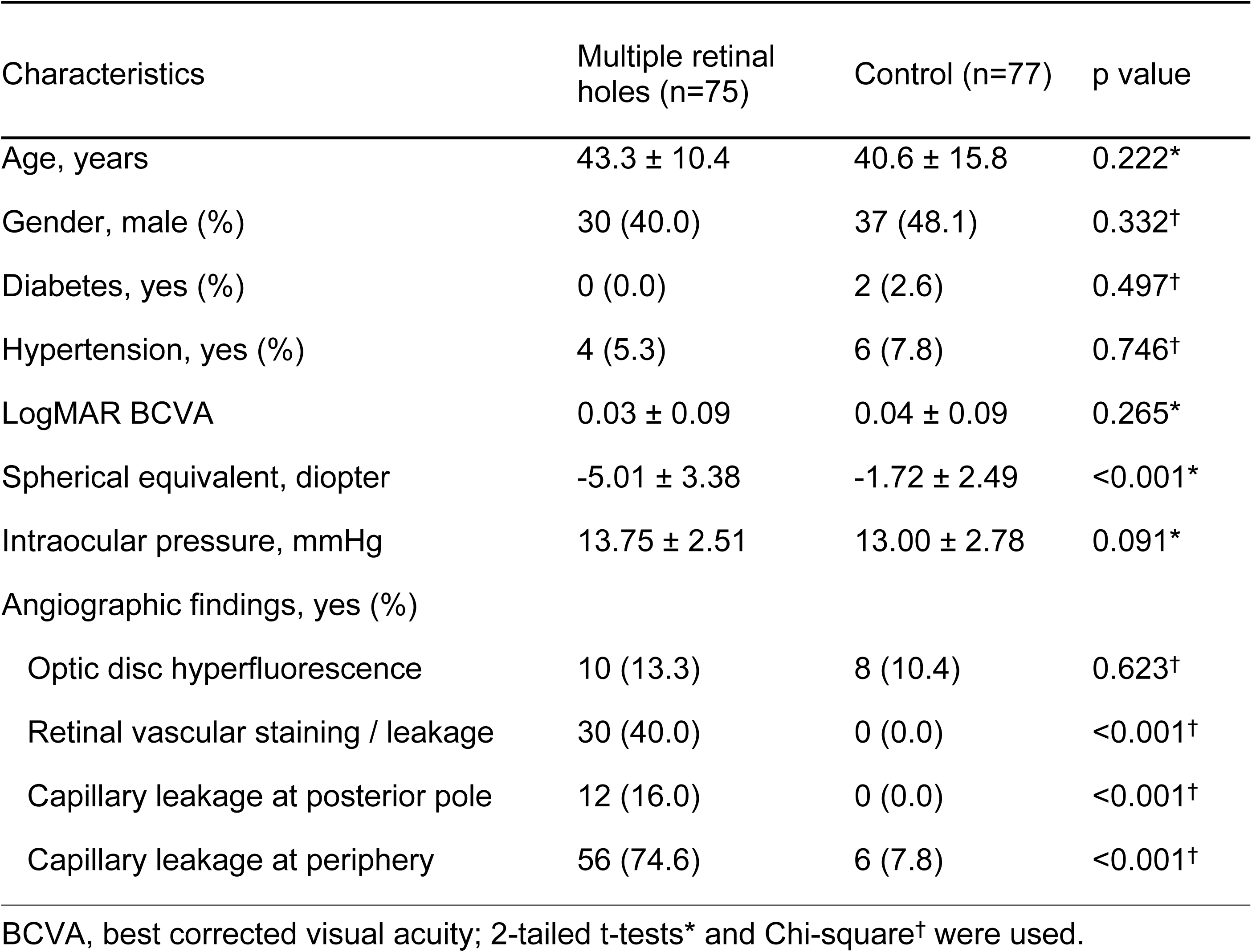
Demographic characteristics and angiographic findings of enrolled patients.

| Characteristics | Multiple retinal holes (n=75) | Control (n=77) | p value |
| --- | --- | --- | --- |
| Age, years | 43.3 ± 10.4 | 40.6 ± 15.8 | 0.222* |
| Gender, male (%) | 30 (40.0) | 37 (48.1) | 0.332<sup>†</sup> |
| Diabetes, yes (%) | 0 (0.0) | 2 (2.6) | 0.497<sup>†</sup> |
| Hypertension, yes (%) | 4 (5.3) | 6 (7.8) | 0.746<sup>†</sup> |
| LogMAR BCVA | 0.03 ± 0.09 | 0.04 ± 0.09 | 0.265* |
| Spherical equivalent, diopter | -5.01 ± 3.38 | -1.72 ± 2.49 | <0.001* |
| Intraocular pressure, mmHg | 13.75 ± 2.51 | 13.00 ± 2.78 | 0.091* |
| Angiographic findings, yes (%) |  |  |  |
| Optic disc hyperfluorescence | 10 (13.3) | 8 (10.4) | 0.623<sup>†</sup> |
| Retinal vascular staining / leakage | 30 (40.0) | 0 (0.0) | <0.001<sup>†</sup> |
| Capillary leakage at posterior pole | 12 (16.0) | 0 (0.0) | <0.001<sup>†</sup> |
| Capillary leakage at periphery | 56 (74.6) | 6 (7.8) | <0.001<sup>†</sup> |

BCVA, best corrected visual acuity; 2-tailed t-tests\* and Chi-square<sup>†</sup> were used.

UWFA detected optic disc hyperfluorescence in 10 of 75 eyes (13.3%) in the case group, and 8 of 77 eyes (10.4%) in the control group (*p*=0.623). Retinal vascular staining/leakage and capillary leakage at the posterior pole were found in 30 eyes (40.0%) and 12 eyes (16.0%), respectively, in the case group; while no eye showed retinal vascular staining/leakage or capillary leakage at the posterior pole in the control group (*p*<0.001, for each). Peripheral capillary leakage was detected in 56 eyes (74.6%) in the case group, and in 6 eyes (7.8%) in the control group (*p*<0.001). No eye showed macular edema, retinal capillary nonperfusion, neovascularization, retinal staining, or subretinal pooling.

To evaluate the factors associated with the severity of angiographic abnormalities in eyes with multiple retinal holes, we performed a regression analysis of ASUWG scores (Table 2). Univariate analysis revealed that the following were associated with a higher ASUWG score: spherical equivalent (r= −0.275, *p*=0.021), and number of retinal holes (r=0.271, *p*=0.027). No differences in ASUWG scores were detected by the two-tailed t-test for gender (*p*=0.929). After adjustments for age, and gender multivariate regression analysis revealed that the spherical equivalent was independently associated with a higher ASUWG score (r^2^=0.161, *p*=0.001; Fig 2).

**Table 2.**
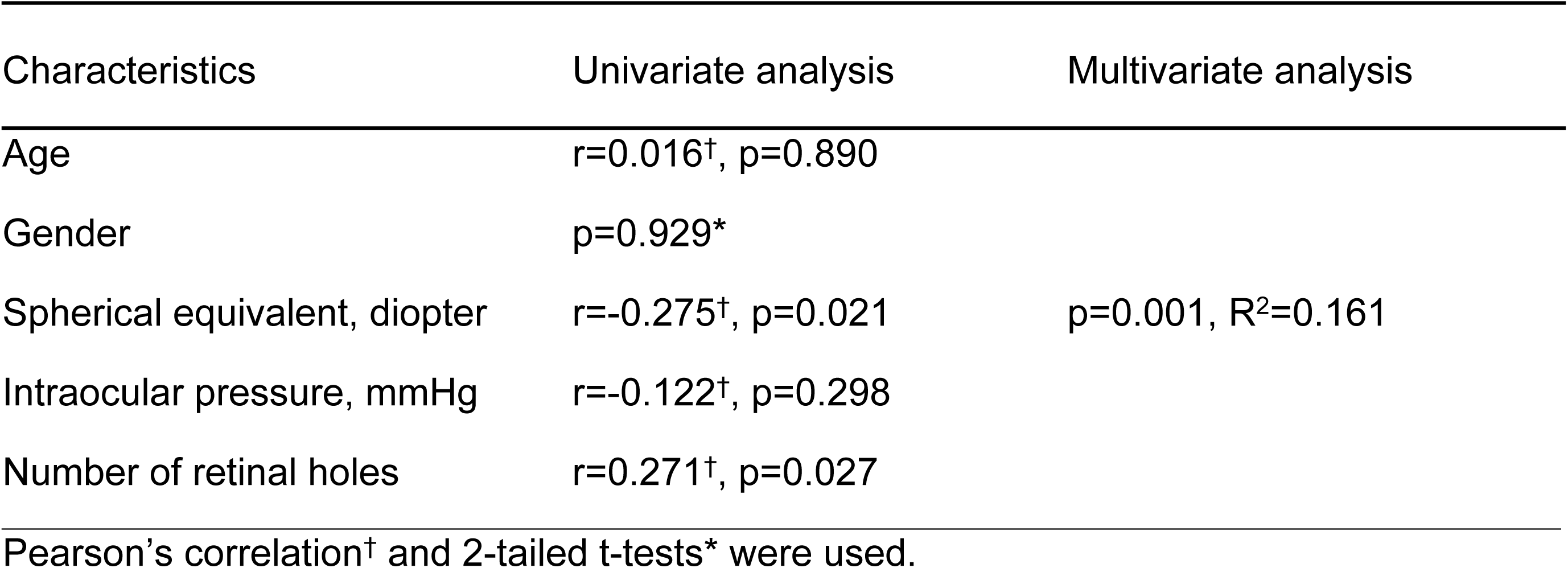
Factors associated with the high ASUWG score in patients with multiple retinal holes.

| Characteristics | Univariate analysis | Multivariate analysis |
| --- | --- | --- |
| Age | $r=0.016^{\dagger}$, $p=0.890$ | |
| Gender | $p=0.929^*$ | |
| Spherical equivalent, diopter | $r=-0.275^{\dagger}$, $p=0.021$ | $p=0.001$, $R^2=0.161$ |
| Intraocular pressure, mmHg | $r=-0.122^{\dagger}$, $p=0.298$ | |
| Number of retinal holes | $r=0.271^{\dagger}$, $p=0.027$ | |

Pearson's correlation<sup>†</sup> and 2-tailed t-tests<sup>\*</sup> were used.

**Figure 2.**
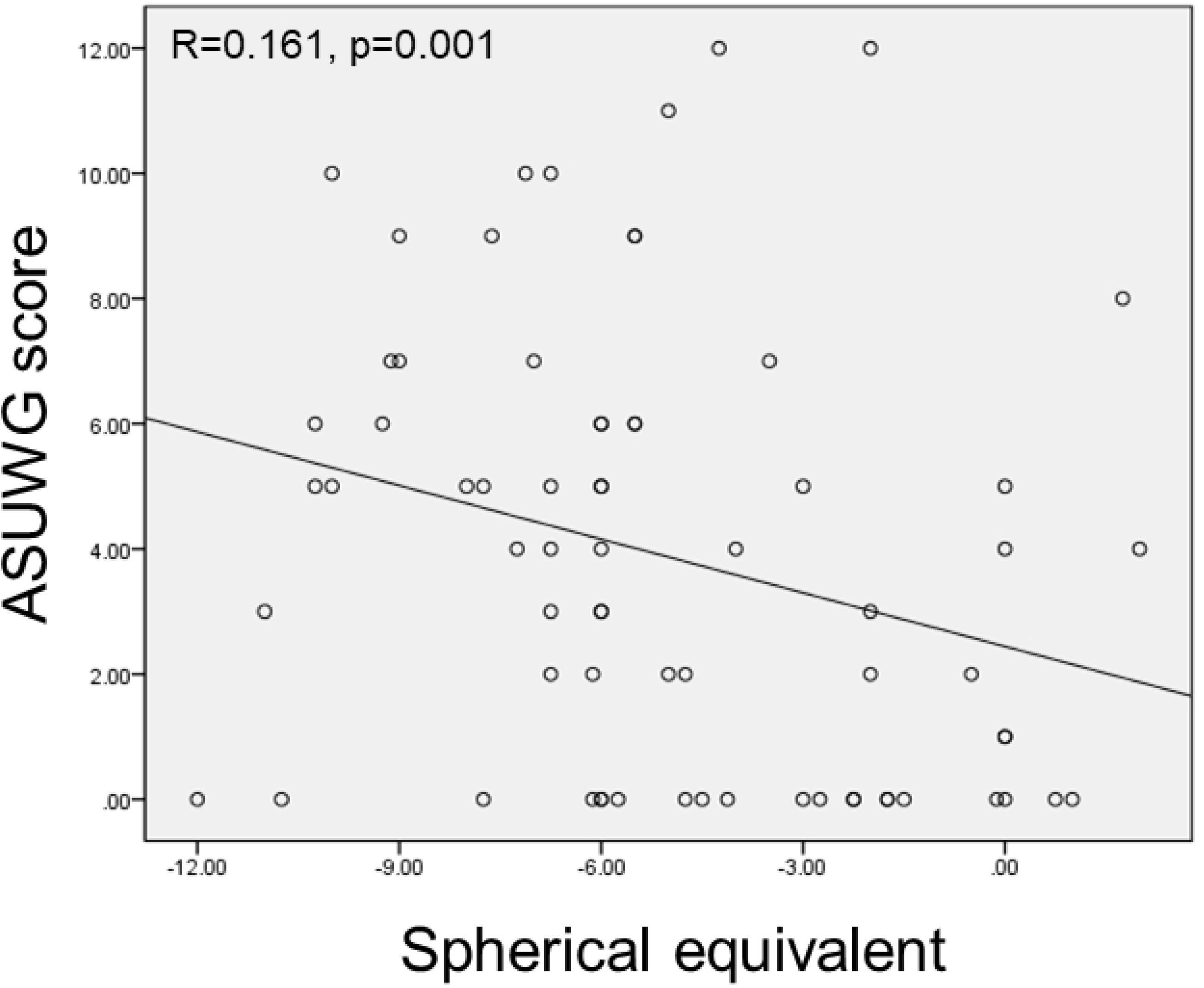
A scatter plot showing the correlation between the refractive error and the Angiography Scoring for Uveitis Working Group (ASUWG) score (r=0.161, *p*=0.001).

Figure 3 shows a representative UWFA image of a patient with multiple retinal holes. The patient was a 21-year old female referred for localized RRD with multiple retinal holes, which were found in the preoperative examination for refractive surgery. A fundus examination revealed a retinal hole-associated localized RRD in the superotemporal quadrant of her right eye, and an atrophic hole without RRD in her left eye, but no vitreous traction on the atrophic holes in either of her eyes. Neither eye showed obvious PVD. In addition, opaque discoloration was detected in the peripheral retina adjacent to the retinal holes in both eyes. OCT image of the discolored retina showed adhesion of vitreous cortex on the peripheral retinal surface adjacent to the retinal hole. UWFA showed perivascular leakages in the peripheral retina, with the extent of the leakages being in accord with the opaque discoloration.

**Figure 3.**
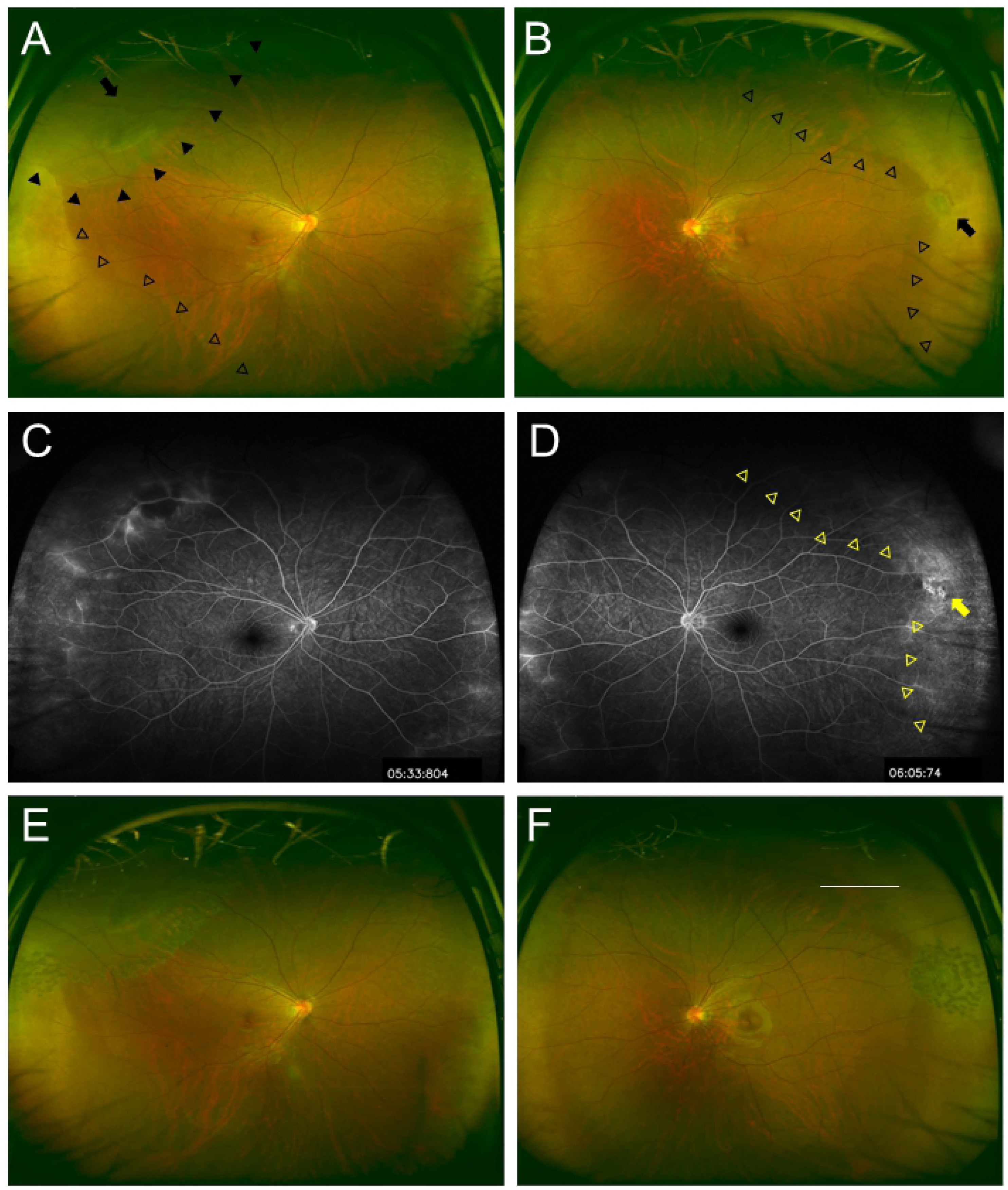
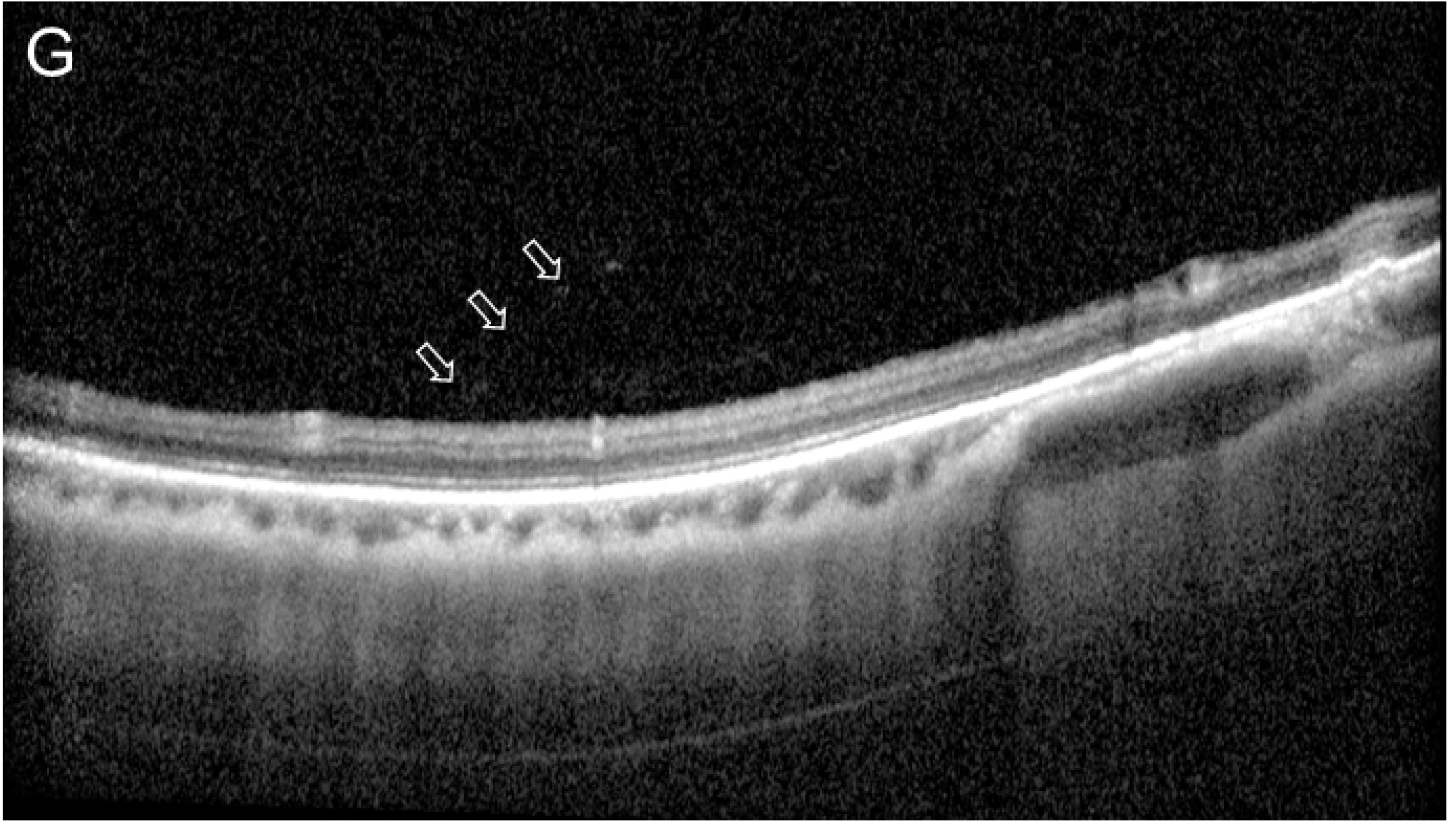
Representative images of an 18-year-old female patient who was referred for asymptomatic RRD in her right eye and bilateral multiple retinal holes which were detected during the screening examination for refractive surgery. (A) The right eye showed focal retinal detachment at the supero-temporal retina with multiple holes (black arrowheads) and discoloration at the infero-temporal retina (blank arrowheads). (B) A retinal hole at temporal retina (arrow) and retinal discoloration (blank arrowheads) was detected in the extension of the retinal hole detected in her left eye. (C and D) UWFA showed bilateral peripheral retinal vascular leakages, and peripheral capillary leakages, which included the area of RRD and the retinal hole. (E and F) Barrier LASER photocoagulation was done to prevent the progression of RRD in both eyes. (G) OCT imaging of the peripheral retina showed adhesion of vitreous cortex (blank arrows) on the peripheral retinal surface adjacent to the retinal hole.

## Discussion

Although RRD is a vision-threatening condition, the pathogenesis of the retinal break, which is the most important determinant factor for RRD, is not fully understood. It is commonly believed that the mechanism involves a vitreoretinal adhesion associated with PVD, which frequently results in retinal tears. However, the pathophysiology of focal retinal thinning or atrophy leading to a retinal hole is largely unknown. We found in this study that most patients with multiple retinal holes showed vascular or capillary leakages at the periphery.

Retinal vascular or capillary leakages can be caused by various retinal pathologic conditions, such as increased venous pressure from vascular occlusion, vascular endothelial cell damage from diabetes,^10^ increased inflammatory cytokines from uveitis,^11^ and mechanical traction from vitreoretinal interface abnormalities.^12^ In our study however, no eye with multiple retinal holes showed retinal vascular occlusion, diabetic retinopathy, or uveitis. We speculate that peripheral vitreoretinal interface abnormalities which are frequently shown in myopic eyes might play a role in the vascular or capillary leakages seen in our patients.

Myopic eyes tend to have vitreoretinal interface abnormalities resulting from the abnormal coordination between the eye’s relatively rigid inner structures and progressively elongated sclera and retina.^13^ Common abnormalities include: epiretinal membranes, retinoschisis, lamellar holes, and paravascular retinal cysts.^14-17^ Scleral elongation can be happen at the posterior pole, as well as equatorially.^18^ We speculate that a patient with posterior pole-type myopia would exhibit vitreoretinal interface abnormalities in the macular area, while equatorial-type myopia would manifest vitreoretinal interface abnormalities in the peripheral retina, as was seen in our patients.

The peripheral vitreoretinal interface abnormalities in myopic eyes have been the subject of a few studies.^19^ Avascular areas and dye leakages from microaneurysms (MA) or telangiectasia were shown to be significantly higher in myopic compared to emmetropic eyes (27% in myopic eyes and 0% in emmetropic eyes), but there was no significant difference in the incidence of capillary telangiectasia and MA. In our study, 74.6 % of eyes with multiple retinal holes showed capillary leakages in the periphery, suggesting an independent relationship between multiple retinal holes and dye leakages. We speculate that chronic traction on the vitreoretinal interface induced by eyeball growth in equatorial lesions might play a role in the peripheral vascular leakages and the formation of retinal holes in myopic patients, as shown in Fig 3.

The development of retinal tear-associated RRD is largely explained by the progression of PVD, since residual attached vitreous cortex can lead to retinal breaks due to abrupt traction. In the general population, PVD typically occurs between the ages of 45 and 65 years.^20^ On the other hand, atrophic retinal hole-associated RRDs tend to be indolent and are found in patients who are younger and more myopic.^21-23^ The presence of PVD can be detected with a dilated fundus examination, and recent OCT technology can provide clear images of the vitreoretinal interface in the posterior pole. However, it is difficult to obtain detailed information about the vitreoretinal interface of peripheral retina in eyes even with clear vitreous. There was no evidence of vitreous detachment on macular area in our patients with multiple retinal holes. We speculate that the partial detachment of vitreous cortex on the peripheral retina induced by equatorial elongation of the eyeball, would explain the development of retinal holes in young myopic eyes with no PVD on the posterior pole. Further study to obtain more detailed information about the temporal changes in the peripheral vitreoretinal interface is warranted to confirm this study.

## Financial Support

“None”

## Conflict of Interest

“No conflicting relationship exists for any author”

